# Improved Quality Metrics for Association and Reproducibility in Chromatin Accessibility Data Using Mutual Information

**DOI:** 10.1101/2023.04.26.538354

**Authors:** Cullen Roth, Vrinda Venu, Vanessa Job, Nicholas Lubbers, Karissa Y. Sanbonmatsu, Christina R. Steadman, Shawn R. Starkenburg

## Abstract

**Background:** Correlation metrics are widely utilized in genomics analysis and often implemented with little regard to assumptions of normality, homoscedasticity, and independence of values. This is especially true when comparing values between replicated sequencing experiments that probe chromatin accessibility, such as assays for transposase-accessible chromatin via sequencing (ATAC-seq). Such data can possess several regions across the human genome with little to no sequencing depth and are thus non-normal with a large portion of zero values. Despite distributed use in the epigenomics field, few studies have evaluated and benchmarked how correlation and association statistics behave across ATAC-seq experiments with known differences or the effects of removing specific outliers from the data. Here, we developed a computational simulation of ATAC-seq data to elucidate the behavior of correlation statistics and to compare their accuracy under set conditions of reproducibility.

**Results:** Using these simulations, we monitored the behavior of several correlation statistics, including the Pearson’s *R* and Spearman’s *ρ* coefficients as well as Kendall’s *τ* and Top-Down correlation. We also test the behavior of association measures, including the coefficient of determination *R*^2^, Kendall’s W, and normalized mutual information. Our experiments reveal an insensitivity of most statistics, including Spearman’s *ρ*, Kendall’s *τ*, and Kendall’s W, to increasing differences between simulated ATAC-seq replicates. The removal of co-zeros (regions lacking mapped sequenced reads) between simulated experiments greatly improves the estimates of correlation and association. After removing co-zeros, the *R*^2^ coefficient and normalized mutual information display the best performance, having a closer one-to-one relationship with the known portion of shared, enhanced loci between simulated replicates. When comparing values between experimental ATAC-seq data using a random forest model, mutual information best predicts ATAC-seq replicate relationships.

**Conclusions:** Collectively, this study demonstrates how measures of correlation and association can behave in epigenomics experiments. We provide improved strategies for quantifying relationships in these increasingly prevalent and important chromatin accessibility assays.

## Background

Epigenetic modifications play an important role in regulating multiple cellular processes ranging from DNA replication to gene expression. These covalent additions to DNA and histone proteins do not alter the underlying DNA sequence, but rather, help modulate chromatin structure resulting in distinctive phenotypes. Genome-wide epigenetic modifications can be determined using several techniques: the gold-standard is chromatin immunoprecipitation followed by sequencing (ChIP-seq) [1, 2, 3]. Chromatin accessibility, or the analysis of the regions that are available for DNA:protein interactions potentially resulting in gene expression, is measured using an enzyme-driven assay called transposase-accessible chromatin via sequencing (ATAC-seq) [4]. These two methods have distinct advantages in probing the state of the epigenome, and both approaches generate paired-end sequencing libraries. These data are mapped to the genome to determine the loci that are occupied with a particular epigenetic modification or the loci that are localized within an open, accessible region. Epigenetic modifications and chromatin accessibility are visualized as peaks resulting from the aggregation of sequencing reads [5]. As such, many software platforms used for analysis of ChIP-seq and ATAC-seq data sets use ‘peak calling’ to determine locations of epigenetic modifications or accessible chromatin regions [6, 7, 8, 9].

To ensure significance and consistency of identified peaks, best practices have been defined for quantifying reproducibility across experimental replicates [8, 10]. These include several quality control metrics and workflows that standardize analysis and enable comparison among different experiments [10]. These standards apply to the total number of sequenced reads, total number of identified significant peaks, and concentration of sequenced reads within said peaks. For example, pseudo-replication was developed for ChIP-seq analysis to assess the amount of variation between biological replicates [8]. In this protocol, synthetic replicates are created from true, experimentally derived data: to do this, aligned reads are merged from two true replicates and randomly reassigned into new alignments to create two synthetic replicates. This permutation practice homogenizes (and splits) signals present within the true, observed replicates, generating the null hypothesis of near perfect correlation between pseudo-replicates. Peak calling is then also conducted on pseudo-replicates, and the read counts of peaks conserved between the two pseudo-replicates are compared to the observed peaks in the true replicates. Landt *et al.* (2012) proposed that experiments, whose number of observed peak counts (among true replicates) divided by the total number of pseudo peaks (between pseudo-replicates), which nears a value of one, are broadly reproducible [8]. The ENCODE project has since extended this practice to ATAC-seq experiments [11, 12].

To better understand experimental reproducibility, many studies also conduct correlation analysis on binned signals between ATAC-seq replicates [13, 14, 15]. In such analyses, for each replicate, the genome is binned into smaller, contiguous regions, for example using windows of ten kilobase pairs [13]. The number of mapped sequenced fragments (defined by a pair of mapped reads) that overlap these bins are counted and standardized to fragments per kilobase pair per million reads (Fpkm) [16]. These Fpkm counts are then compared between replicates using correlation and association statistics such as Pearson’s *R* or the coefficient of determination (*R*^2^), respectively. Values from these statistics trending toward a value of one generally indicate a reproducible experiment [17].

Correlation analysis is a useful tool, not singularly purposed for the analysis of reproducibility in ATAC-seq experiments. Such analysis can be found within studies of chromosome accessibility in cancer, ageing of human stem cells, cellular diversity, or new ATAC-seq protocols [18, 19, 20, 21, 22, 23]. Furthermore, correlation analyses are ubiquitous, found in the fields of genetics, RNA-seq experiments, and in studies of 3D chromatin architecture [24, 25, 26, 27, 28, 29, 30, 16]. Given their popularity and use in genomic and epigenetic studies, software suites—for example deeptools and HiCExplorer—have developed methods and tools for calculating correlation metrics between replicates and experiments [13, 31, 32, 33, 34].

The natural properties of data from genomic and epigenomic experiments make the application of com-monly used correlation and association statistics, for example Pearson’s *R* and *R*^2^, potentially problematic as none of these data (ATAC-, ChIP-, or Hi-C seq) are normally distributed [35]. Both ATAC-and ChIP-seq experiments are defined by numerous, loci-specific peaks of signal generated by the accumulation of sequencing reads [3, 4]. Mapped sequenced fragments may overlap contiguous genomic bins used in analysis, producing non-independent data points [24]. Conversely, regions lacking assayed modifications or with inaccessible chromatin will have little to zero signal for ChIP-seq or ATAC-seq data, respectively. Furthermore, during correlation analysis, several genomic bins may overlap an inaccessible chromatin region that is reproducible, appearing in both the ATAC-seq replicates (or experiments) being compared. As such, each of these bins will acquire zero Fpkm and within the bi-variate distribution formed between the replicates. These data points, which appear as zero Fpkm in both replicates, are referred to here as co-zeros. Some analysis programs, like deeptools, HiCExplorer, and HiCcompare, offer options to remove co-zeros prior to analysis [31, 34, 29]. However, there is no published guidance on this practice, and while the co-zero values are a feature common across genomic and epigenomic data sets [36], the effect of removing such features on correlation statistics has not been explored. Despite the known features of genomic and epigenomic data, and the underlying assumptions of statistical tests, there have been few studies that explore their expected behavior, accuracy, and use of alternative statistics determining reproducibility of such data [26, 27].

Here, we present a computational approach to generate synthetic ATAC-seq replicates to explore the behavior of various correlation and association metrics for epigenomics datasets. These synthetic ATAC-seq replicates are generated from eight true data sets to capture features uniquely present within ATAC-seq experiments. We have developed a random subsampling strategy to generate synthetic replicates with varying portions of shared peaks, as a proxy for reproducibility. Across our simulations, we apply the Pearson’s *R* [37, 38, 39] and Spearman’s *ρ* [40] and monitor their behavior, including the effect of removing co-zeros. Additionally, we demonstrate the behavior of other statistics, including non-parametrics such as Kendall’s *τ* [41, 42, 43, 44] and an information theoretic approach, normalized mutual information [45, 46], to determine their utility in assessing epigenomics data. Finally, we build a random forest model [47] using the normalized mutual information and *R*^2^ coefficient between experiments to predict the biological relationships between replicates. Overall, our results demonstrate an improvement in the expected behavior of all statistics after removing co-zeros and normalized mutual information emerges as a promising statistic for measuring association between ATAC-seq samples.

## Results

### ATAC-seq data characteristics and subsamping strategy for synthetic replicate generation

To study the behavior of correlation measurements between ATAC-seq replicates, we analyzed data from three experiments using the A549, human lung cell line and implemented a subsampling paradigm to generate synthetic replicates. Across these experiments, the total number of reads mapped to the human reference genome varied from 15 million to nearly 43 million (Table 1). The number of genome-wide peaks found in the ATAC-seq samples varied across experiments and between replicates, ranging from approximately 80 to 130 thousand (Table 1). The fraction of sequenced read-pairs mapped in peaks (i.e. the FrIP score as defined by the ENCODE project [8, 11]), was greater than 0.34 for all of the A549 ATAC-seq samples (Table 1). These samples displayed high spatial correlation of peaks across replicates (Figure 1A). Counting all whole fragments per kilobase per million (WFpkm), every ten kilobases, we observed a high statistical correlation between replicates, with average Pearson’s *R* of 0.86, 0.87, and 0.94 (*p*-values *<* 0.05) between the technical replicates of the three biological replicate experiments (Figure 1B).

**Figure 1:**
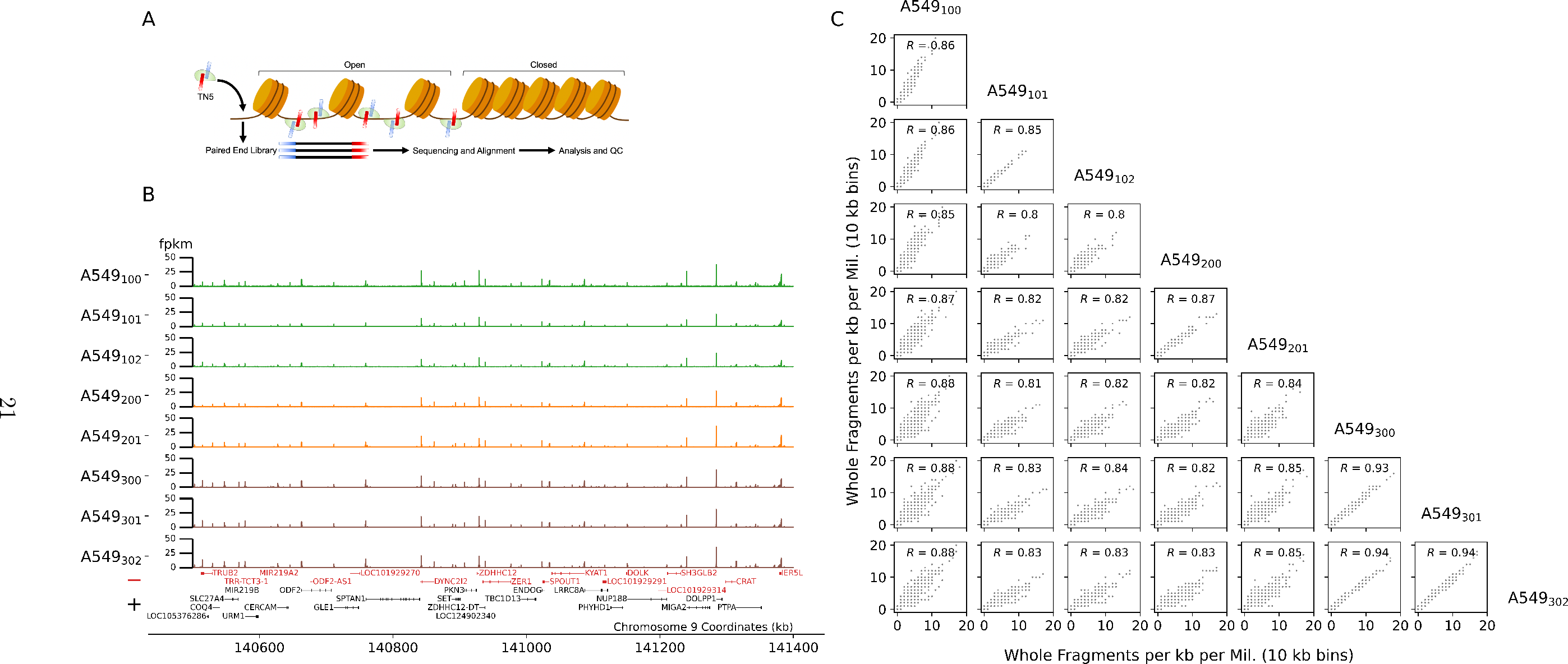
ATAC-seq profiles of chromosome 9 form A549 cells. **A**: TN5 binds to open chromatin, cutting DNA and adding primers to generate a paired-end sequencing library. **B**: A549, ATAC-seq replicates along chromosome 9. Samples were generated using fresh cells (green) and previously cryo-preserved cell cultures (orange and brown). Positively (black) and negatively oriented genes are annotated along the bottom. **C**: Pair-wise, bi-variate scatter plots of whole fragments per kb per million values (x- and y-axis) using 10 kb genomic bins between A549, ATAC-seq replicates. Sample names are annotated along the diagonal. Pair-wise Pearson’s correlation statistic is annotated within subplots.

**Table 1:** ATAC-seq Experiments Used, Mapped Reads, Peak Counts and FrIP Scores

| Sample Title | Cell Line | Mapped Reads | MACS2 Peaks | FrIP | Source |
| --- | --- | --- | --- | --- | --- |
| A549<sub>000</sub> | A549 | 259029456 | 201532 | 0.5898 | ENCSR032RGS |
| A549<sub>001</sub> | A549 | 329679445 | 194975 | 0.5994 | ENCSR032RGS |
| A549<sub>002</sub> | A549 | 211291691 | 206536 | 0.5596 | ENCSR032RGS |
| A549<sub>100</sub> | A549 | 23987725 | 110323 | 0.588 | This study |
| A549<sub>101</sub> | A549 | 22605005 | 81917 | 0.3404 | This study |
| A549<sub>102</sub> | A549 | 17618743 | 82496 | 0.3702 | This study |
| A549<sub>200</sub> | A549 | 35069198 | 90386 | 0.3515 | This study |
| A549<sub>201</sub> | A549 | 15377297 | 79933 | 0.4202 | This study |
| A549<sub>300</sub> | A549 | 42567716 | 130475 | 0.636 | This study |
| A549<sub>301</sub> | A549 | 28744542 | 107737 | 0.6391 | This study |
| A549<sub>302</sub> | A549 | 35836016 | 117087 | 0.6595 | This study |
| GM12878<sub>400</sub> | GM12878 | 46889870 | 114746 | 0.7159 | ENCSR095QNB |
| GM12878<sub>401</sub> | GM12878 | 49588811 | 134743 | 0.6452 | ENCSR095QNB |
| HepG2<sub>500</sub> | HepG2 | 48113686 | 173756 | 0.4257 | ENCSR042AWH |
| HepG2<sub>501</sub> | HepG2 | 48246610 | 135767 | 0.4605 | ENCSR042AWH |
| IMR-90<sub>600</sub> | IMR-90 | 47543633 | 178156 | 0.5363 | ENCSR200OML |
| IMR-90<sub>601</sub> | IMR-90 | 61359070 | 200216 | 0.6104 | ENCSR200OML |
| K562<sub>700</sub> | K562 | 48217636 | 178230 | 0.5112 | ENCSR483RKN |
| K562<sub>701</sub> | K562 | 52270533 | 176789 | 0.5196 | ENCSR483RKN |
| RWPE2<sub>800</sub> | RWPE2 | 55152003 | 166239 | 0.474 | ENCSR080SNF |
| RWPE2<sub>801</sub> | RWPE2 | 43166947 | 177496 | 0.4555 | ENCSR080SNF |
| RWPE2<sub>802</sub> | RWPE2 | 48162285 | 154758 | 0.4652 | ENCSR080SNF |
| WTC11<sub>900</sub> | WTC11 | 74558506 | 245677 | 0.5505 | ENCSR541KFY |
| WTC11<sub>901</sub> | WTC11 | 79335328 | 277824 | 0.5732 | ENCSR541KFY |

For simulations, synthetic replicates were generated using the paired-end read alignment profiles from the eight ATAC-seq samples we generated. For each simulation, two synthetic replicates were initiated by duplicating a given true ATAC-seq experiment (Figure 2A). Within the true ATAC-seq data set, reproducible, significant peaks were identified (see Methods). From these, a random portion of peaks was chosen to vary between the two synthetic replicates. This was accomplished by subsampling 85% of the aligned sequenced fragments within each of the randomly chosen peaks between the two synthetic replicates (Figure 2B and 2C). This process was repeated, randomly varying the common peaks from 1 to 95% of peaks between the two synthetic replicates. Finally, across all simulations, for each pair of synthetic replicates, the WFpkm values were calculated in ten kilobase windows and used in statistical comparisons (Figure 3A).

**Figure 2:**
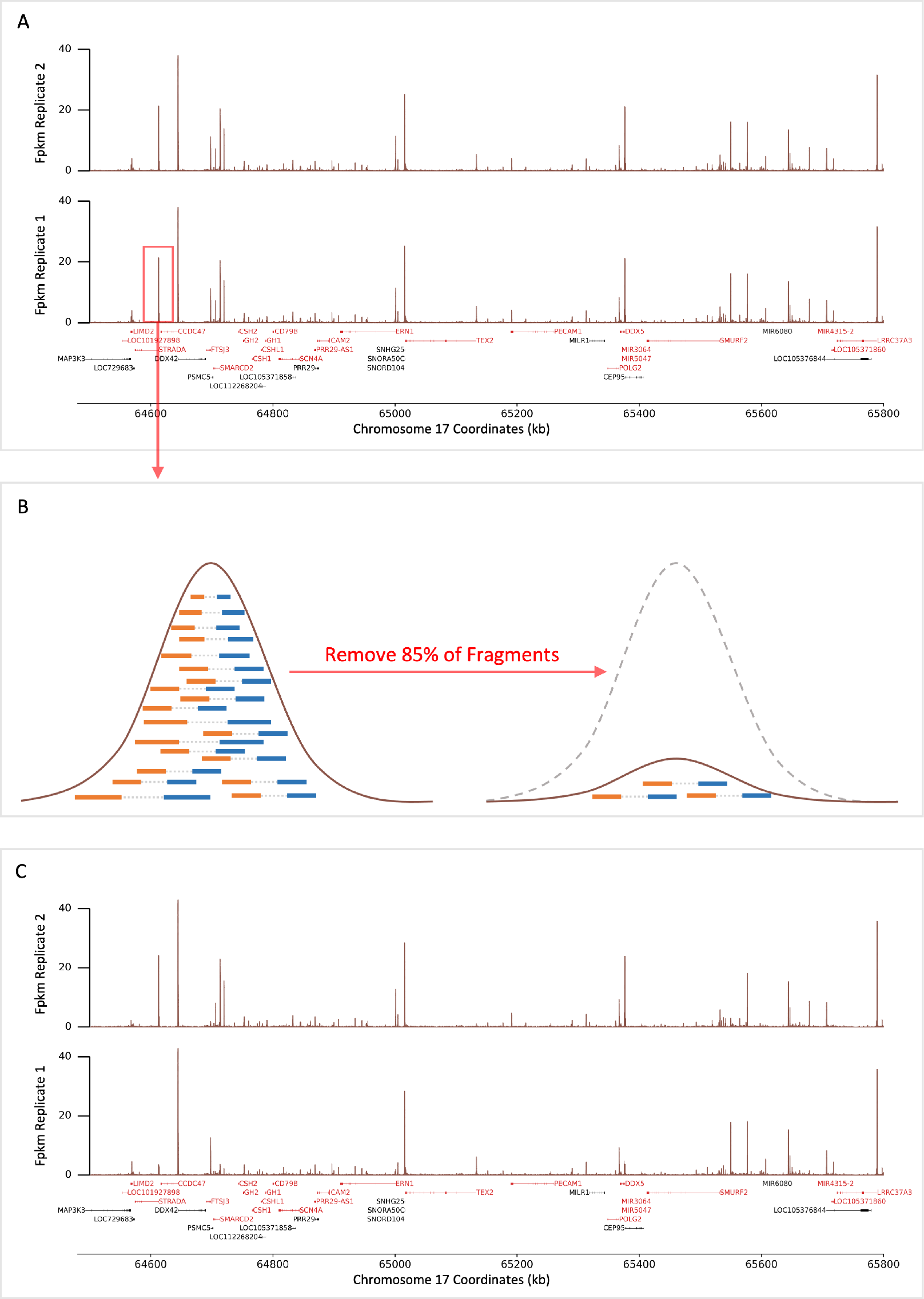
Synthetic replicate generation via peak down-sampling. **A**: An example region along chromosome 17 of true, A549 ATAC-seq data. Real ATAC-seq signal (brown lines) is used to initialize two synthetic replicates. Red and black horizontal bodies depict negatively and positively oriented genes, respectively. **B**: A portion of the genome-wide significant peaks (ranging from 0 - 1) are chosen randomly between the two synthetic replicates. Within one of the replicates, 85% of paired reads (blue and orange rectangles connected by grey dotted line) are removed to down-sample signal within that locus. **C**: Example of two synthetic replicates with a known portion of peaks varying between them.

**Figure 3:**
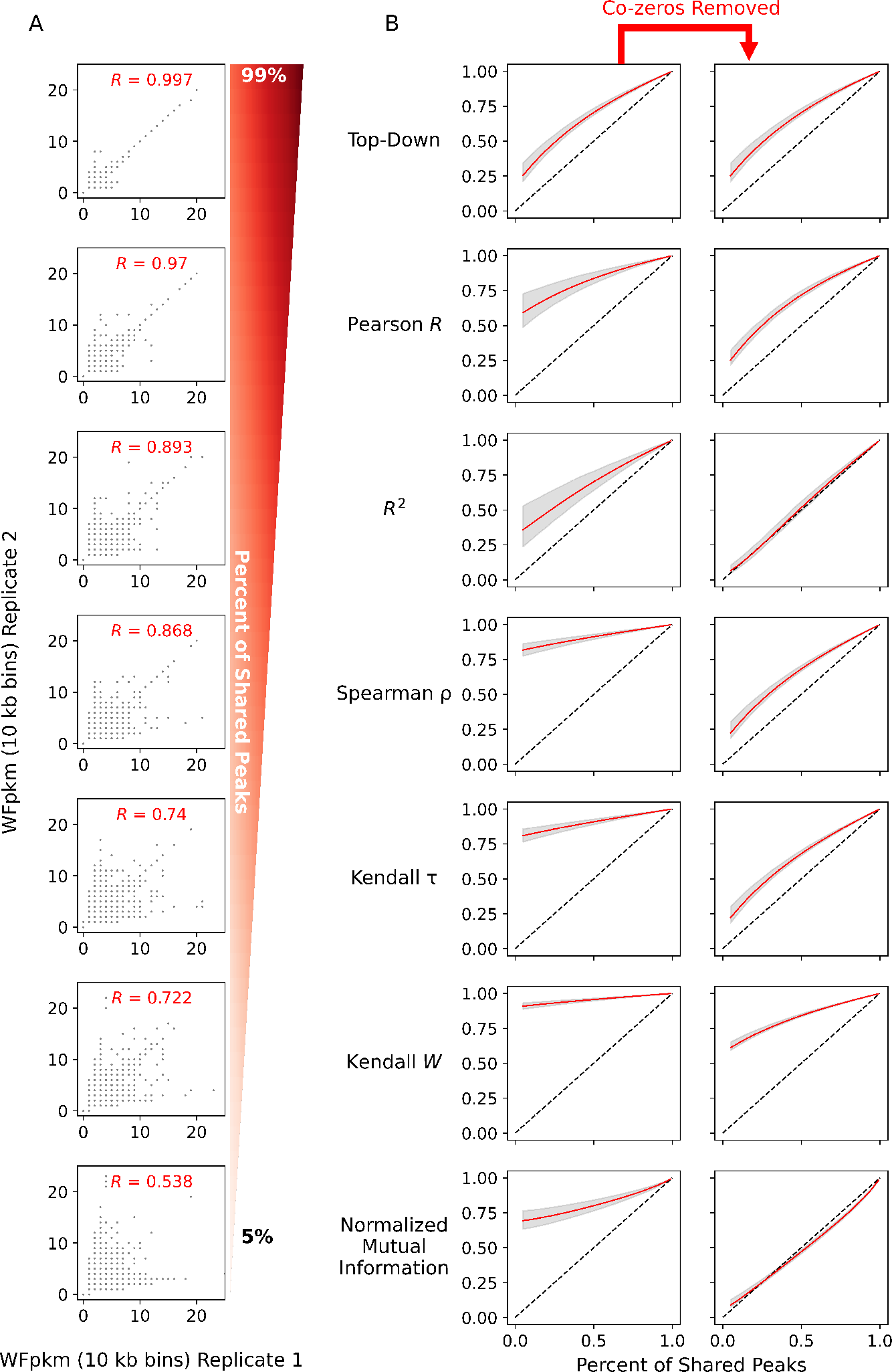
Synthetic replicate bivariate plots and statistical profiles. **A**: Scatter plots displaying counts per genomic bin (10 kb in size) of whole fragments per kilobase per million between two synthetic replicates (x- and y-axis) generated in process Figure 2A – C. The percentage of shared peaks decreases between the two simulated replicates from top to bottom. **B**: Correlation and association values (y-axis) as a function of percentage of shared peaks between synthetic replicates (x-axis). Red and grey curves depict the mean and 95% CI (respectively) values across simulations. A grey, dashed line marks a one-to-one relationship between the x- and y-axis. Left and right columns display change in values as a function of removing co-zeros.

### Top-down correlation displays best behavior in correlation analysis across simulations

Across these down sampling simulations, correlation and association statistics were calculated between each pair of synthetic replicates. The Wfpkm counts were used between synthetic replicates in statistical analysis (Figure 3A). The values of correlation and association statistics were calculated across simulations, as a function of the number of shared peaks between synthetic replicates (Figure 3B) and for each statistic, and the area under the curve (AUC) was used in comparisons (Supplementary Figure S1). Of the correlation statistics, the Top-Down correlation statistic had the smallest average AUC of 0.6881 (95% CI: 0.6860 – 0.6906) and was significantly smaller than the average AUC of the Pearson’s *R*, at 0.8284 (95% CI: 0.8237 – 0.8335, *p*-value = 0, bootstrapped difference of mean AUC). Both the two non-parametric statistics, Spearman’s *ρ* and Kendall *τ*, had significantly larger average AUCs compared against the Pearson’s *R* (*p*-values = 0, bootstrapped difference of mean AUC). However, they demonstrated nearly identical AUC profiles compared to each other, with average AUC of 0.9140 (95% CI: 0.9118 – 0.9162) and 0.9096 (95% CI: 0.9074 – 0.9120) respectively (*p*-value = 0.037, bootstrapped difference of mean AUC).

Across the metrics of association, Kendall’s W, normalized mutual information, and the *R*^2^ coefficient, between replicates, the *R*^2^ coefficient exhibited the greatest sensitivity to the change in portion of shared peaks between synthetic replicates (Figure 3B). Across simulations, the average AUC of the *R*^2^ coefficient was 0.7026 (95% CI: 0.6951 – 0.7102). This average AUC was significantly smaller than the average AUC of the Kendall’s W and normalized mutual information, with values of 0.957 (95% CI: 0.9559 – 0.9581) and 0.8197 (95% CI: 0.8153 – 0.8241), respectively (*p*-value = 0, bootstrapped difference of mean AUC).

### Removal of co-zeros improves estimates of correlation and associations

Using this simulation paradigm, we evaluated the efficacy of removing co-zeros from the analysis to determine the impact on correlation and association statistics. Co-zero values were defined as value counts in ATAC-seq experiments that appeared to have zero aligned fragments in a genomic bin of ten kilobases between two replicates (Figure 3B, Supplementary Figure S2). On average, these values can make up nearly 5% of a given bi-variate distribution formed between real ATAC-seq replicates (Supplementary Figure S3). Across all the correlation and association statistics examined here—except for Top-Down correlation—removing the co-zero values significantly reduced the average AUC (Table 2, Figure 3B, Supplementary Figure S1). This finding was unexpected, as co-zeros are a modest portion of the bi-variate distribution formed between two replicates and reproducible data points within the replicates.

**Table 2:** Mean Area Under the Curve Across Simulations

| Statistic | Mean (95% CI) | Mean (95% CI) – Co-zeros removed | $p$-value <sup>a</sup> | $\sigma^2$ <sup>b</sup> |
| --- | --- | --- | --- | --- |
| Top-Down Correlation | 0.6881 (0.6860 – 0.6906) | 0.6872 (0.6850 – 0.6895) | 0.635 | - |
| Pearson $R$ | 0.8284 (0.8237 – 0.8335) | 0.6965 (0.6946 – 0.6984) | 0.0 | 0.0201 |
| $R^2$ | 0.7026 (0.6951 – 0.7102) | 0.5346 (0.5324 – 0.5368) | 0.0 | 0.0329 |
| Spearman $\rho$ | 0.9140 (0.9118 – 0.9162) | 0.6686 (0.6665 – 0.6705) | 0.0 | 0.0136 |
| Kendall $\tau$ | 0.9096 (0.9074 – 0.9120) | 0.6673 (0.6654 – 0.6691) | 0.0 | - |
| Kendall W | 0.9570 (0.9559 – 0.9581) | 0.8343 (0.8333 – 0.8353) | 0.0 | - |
| Normalized Mutual Information | 0.8197 (0.8153 – 0.8241) | 0.5055 (0.5045 – 0.5065) | 0.0 | 0.016 |

<sup>a</sup> The $p$ -value represents the test of differences in mean AUC after removal of co-zeros.
<sup>b</sup> Variation values were calculated during analysis of data from true ATAC-seq experiments.

After removing co-zeros, all the correlation statistics, Top-Down correlation, Pearson’s *R*, Spearman’s *ρ*, and Kendall’s *τ*, displayed nearly identical sensitivity to the change in shared peaks between replicates across simulations (Figure 3B). However, the Pearson’s *R* had the largest average AUC of 0.6965 (95% CI: 0.6946 – 0.6984) followed by the Top-Down statistic (AUC of 0.6872, 95% CI: 0.685 – 0.6895, *p*-value = 0, bootstrapped difference of mean AUC). The Spearman’s *ρ* (mean AUC: 0.6686, 95% CI: 0.6665 – 0.6705) and Kendall’s *τ* (mean AUC: 0.6673, 95% CI: 0.6654 – 0.6691) statistics had the smallest and identical average AUC after removing co-zeros (*p*-value = 0.208, bootstrapped difference of mean AUC). Furthermore, the AUC of the Top-Down correlation statistic was unaltered by the exclusion of co-zero values between synthetic replicates (Figure 3B, Supplementary Figure S1, Table 2, *p*-value = 0.635, bootstrapped difference of mean AUC). This observation was not surprising given how Top-Down correlation places emphasis on larger values, down-weighting smaller values, such as co-zeros [48].

### Normalized mutual information best estimates difference between replicates

Removing co-zero values had a similar effect on association metrics, attenuating and improving the average AUC across the portion of shared peaks between synthetic replicates (Figure 3B, Supplementary Figure S1). Apart from Kendall’s W, the *R*^2^ coefficient and normalized mutual information, on average, displayed a nearly one-to-one relationship with the portion of shared peaks between replicates (Figure 3B). The average AUC of normalized mutual information was 0.5055 (95% CI: 0.5045 – 0.5065) and was smaller than the average AUC of the *R*^2^ coefficient, with a value of 0.5346 (95% CI: 0.5324 – 0.5368, *p*-value = 0, bootstrapped difference of mean AUC). This difference in average AUC indicates that normalized mutual information better follows the designed proportion of shared peaks between synthetic replicates across our simulations, compared to the *R*^2^ coefficient.

As introduced earlier, one parameter in this simulation is the removal of a percentage of aligned read-pairs from within randomly selected peaks (Figure 2B). Initially set at 85%, this parameter was altered to simulate ATAC-seq replicates that are nearly reproducible (at 50%) at every selected peak or broadly unreproducible (at 95%) across all selected peaks. Comparing the results between the two simulation sets with 85 and 95% of reads removed, we observed no significant difference between the two simulations (see Supplementary Data). This is somewhat expected when considering the small difference in magnitude between removing 85 and 95% of reads from within peaks. In simulations with only 50% of read pairs removed from selected peaks, after removing co-zeros, the two statistics that showed the largest response in our simulation were the *R*^2^ coefficient and normalized mutual information (see Additional File 1 and Additional File 2).

### Validation of mutual information analysis on true ATAC-seq data

After its successful implementation on simulated replicates, we next examined how normalized mutual information behaves when used on replicates from true ATAC-seq experiments. For this analysis, additional ATAC-seq experiments were downloaded from the ENCODE project public repository [11]. These included additional replicates of the A549 cell line, as well as ATAC-seq experiments in the HepG2, RWPE2, GM12878, IMR-90, K562, and WTC11 cell lines (Table 1). With this expanded dataset, the Pearson’s *R*, the Spearman’s *ρ*, *R*^2^ coefficient, and normalized mutual information were calculated between all pairs of replicates, with co-zeros removed from analysis (Figure 4). Removing co-zeros reduced the estimates of correlation and association across samples by approximately 0.18, 0.32, 0.23, and 0.30 on average for the Pearson’s *R*, the Spearman’s *ρ*, *R*^2^ coefficient, and normalized mutual information, respectively (Supplementary Figure S4A). These differences were significantly greater than zero (*p*-value *<* 10*^−^*10, Wilcoxon signed-rank test, Supplementary Figure S4B). Between comparisons of true experiments, we observed a co-linear relationship in the values of the normalized mutual information scores and *R*^2^ coefficients (Figure 5A, Pearson’s *R* = 0.96, *p*-value *<* 1*^−^*^10^). Of the four statistics, the *R*^2^ coefficient displayed the largest variation (*σ*^2^ = 0.0329) between true replicates (Table 2).

**Figure 4:**
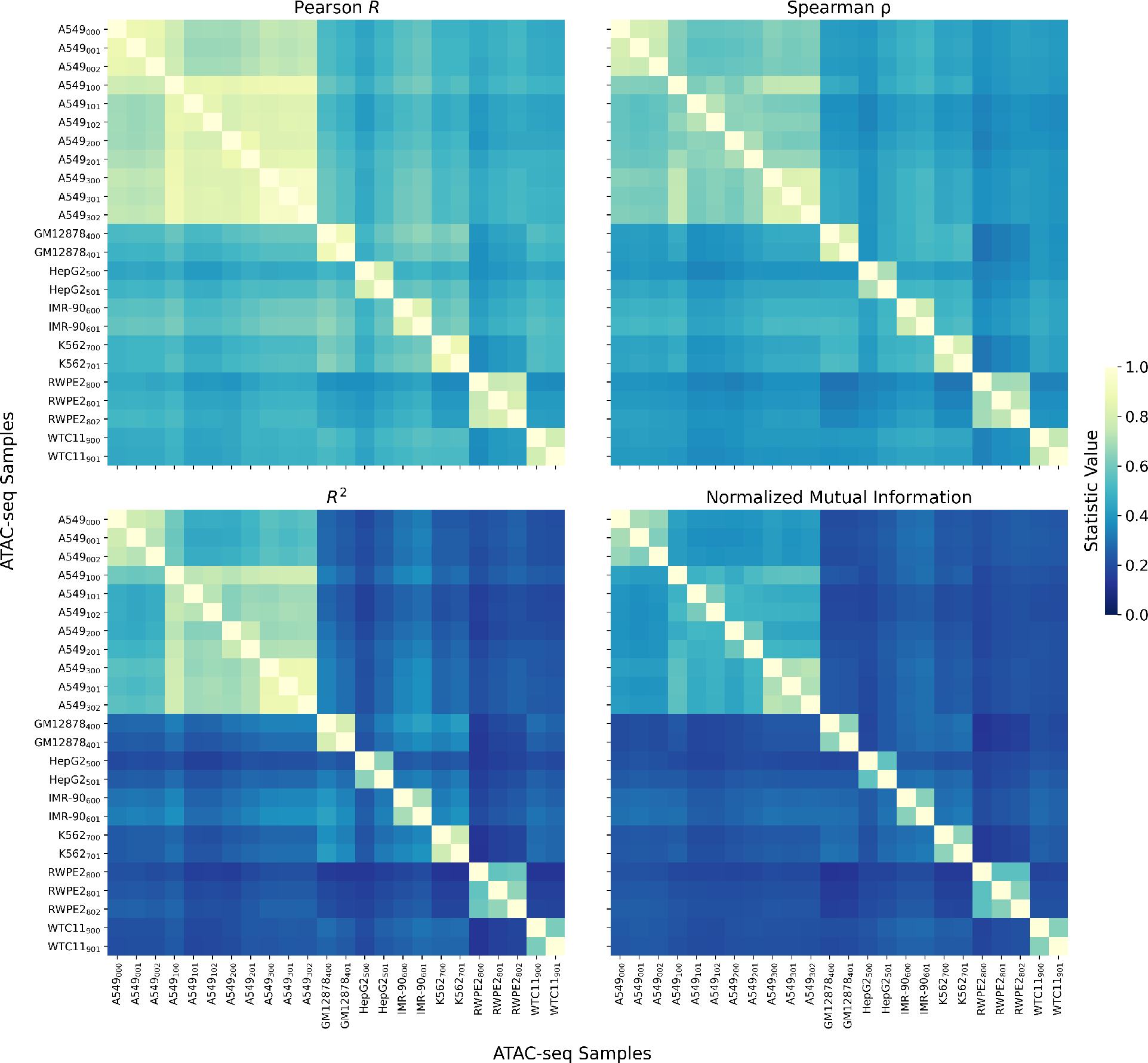
Correlation and association statistics across ATAC-seq experiments. From top-to-bottom, left-to-right, the Person’s *R*, Spearman’s *ρ*, *R*^2^ coefficient, and normalized mutual information across ATAC-seq replicates from the ENCODE project and ATAC-seq experiments on A549 cells generated in this study.

**Figure 5:**
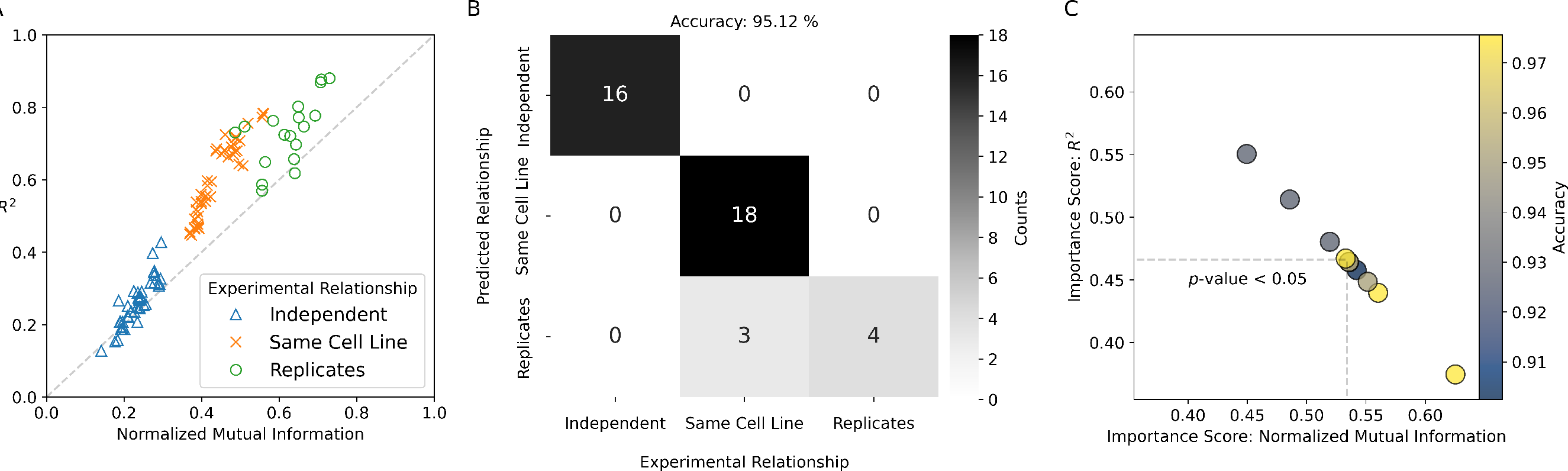
Random forest prediction of experimental relationships. **A**: The coefficient of determination (*R*^2^) versus the normalized mutual information (y- and x-axis, respectively) calculated on binned counts of WFpkm between ATAC-seq experiments. Blue triangles, orange Xs, and green circles mark comparisons between independent experiments, independent experiments using the same cell line, and true experimental replicates, respectively. **B**: Example confusion matrix from a random forest model using *R*^2^ and normalized mutual information as features to predict experimental relationships (y-axis) presented in **A** (x-axis). The confusion matrix depicts results of model on a hold-out set (40% of data, accuracy = 95.12%). Light to dark colors depict the number of counts per class. **C**: Bi-variate plot displaying the change of paired importance scores from ten-fold cross validation between the normalized mutual information (x-axis) and *R*^2^ (y-axis) features. Dashed lines depict the uni-variate means of the normalized mutual information and *R*^2^ scores. Blue and yellow colors depict the level of accuracy for each fold.

### Predicting replicate relationships using normalized mutual information

Given the comparable behavior between normalized mutual information and the *R*^2^ coefficient on true ATAC-seq replicates, we assessed their usefulness in predicting the relationships between experiments. To do this, we utilized a random forest model, using the values of the *R*^2^ coefficient and normalized mutual information between true ATAC-seq experiments as features. Comparisons between any two ATAC-seq experiments (either those from the ENCODE project or generated here) were classified into one of three discrete classes: (1) between independent ATAC-seq experiments in different cell lines, (2) independent experiments using the same cell line, and (3) between true replicates. Plotting the normalized mutual information against the *R*^2^ coefficient calculated between ATAC-seq experiments with the above classifications revealed clustering of experimental relationships between replicates (Figure 5A).

To build our random forest model, we utilized ten-fold cross validation, stratifying on the replicate class. An example confusion matrix from one of these folds demonstrates the model had difficulty distinguishing between independent experiments using the same cell line and true, experimental replicates (Figure 5B). This difficulty also manifested as lower f1-scores and recall for this class (Supplementary Figure S5). The accuracy across these folds ranged from 88 to 98% (Figure 5C). Across the folds, the feature importance score of the *R*^2^ coefficient was inverted with that of normalized mutual information (Figure 5C). Overall, we observed a greater feature importance score for normalized mutual information, with a significant average pair-wise difference between the *R*^2^ coefficient and normalized mutual information of 6.78% (*p*-value *<* 0.05, Wilcoxon signed-rank test).

## Discussion

To improve the assessment of reproducibility in epigenomic data sets, we sought to investigate the use of several correlation and association statistics on binned genomic signals. Our findings suggest that best practices should include analyzing association between compared replicates (or experiments) via normalized mutual information with binned, Fpkm counts rounded to the nearest whole integer, after the removal of co-zero values as input. In choosing a correlation statistic, after removing co-zero values, our results indicate little difference in the outputs from the Pearson’s *R*, Spearman’s *ρ*, Kendall’s *τ*, or Top-Down correlation statistics. Notably, from simulations, we observed that the Top-Down correlation statistic was unaffected by the removal of co-zeros values. As such, this statistic should serve as an alternative for investigators if binned co-zeros values between replicates are retained.

As part of this study, we generated highly correlated, new ATAC-seq experimental replicates of the A549 cell line. Our data highly correlates with previously published ATAC-seq data of the A549 cell line generated by the ENCODE project. Using these data, we generated a novel simulation that utilizes down sampling to generate replicates with known varying signals. While similar simulation studies have been conducted on Hi-C sequencing data [30], to our knowledge, no prior study has examined the behavior of statistical metrics on ATAC-seq data. That said, there are several statistics and methodologies that may be used to analyze this data type, such as Poisson regression [49]. Improving on this simulation design could help generate a framework that allows researchers to develop new statistical tools for hypothesis testing.

In our simulations, we observed that most statistics overestimate the correlation of signal between replicates. One specific strategy we investigated to reduce this inflation was the removal of co-zeros, which is an option present in several bioinformatic software suites [31, 34, 29]. Our analysis demonstrated that removal of these values can provide a more accurate estimate of correlation between replicates as measured by the known number of peaks between replicates. Interestingly, we never observed a correlation value that perfectly trends with the designed number of peaks between synthetic replicates. We also did not observer negative correlation values between the replicate Fpkm counts. The first of these observations can be explained by background autocorrelation still present within our synthetic replicates. The second of these observations may point to a limitation in the design of our simulation, as negative correlation values have been observed in true ATAC-seq profiles [31, 20]

In epigenomics and chromatin accessibility data sets, biological interpretation of the data is dependent upon visualization of “peaks” where accumulation of sequenced reads denotes the presence of a modification or an accessible region. Regions with zero (or nearly zero) aligned sequenced reads are deemed unmodified or inaccessible and largely ignored when interpreting data. Correlation statistics should provide biologists with the confidence that replicates are truly comparable. As stated above, the inclusion of co-zeros seems to inflate values of most correlation and association statistics. Thus, removal of co-zeros formed by the genomic bins that overlap and account for inaccessible regions may be warranted.

Using our simulation, we also examined the behavior of three association statistics, which we distinguish from the set of correlation statistics as those metrics ranging in value from zero to one. These association statistics were the *R*^2^ coefficient, normalized mutual information statistic, and Kendall’s W. Prior to the removal of co-zeros, the only association statistic that displayed any sensitivity to the change in shared peaks between replicates was the *R*^2^ coefficient. Co-zeros inflate the value of this statistic by reducing the total summed error between data points during calculation. Similarly, co-zeros increase the information gained between replicates when calculating the normalized mutual information score. In other words, knowing a replicate has a value of zero at a given genomic bin provides information that there is a zero at the corresponding bin within the other replicate. After removing co-zeros, we saw a large improvement in the sensitivity of both these statistics.

Curiously, Kendall’s W displayed the least sensitivity to the designed peak counts between synthetic replicates. This statistic was of interest given Kendall’s W is capable of simultaneously examining the ranks of more than two input samples [50, 41]. This would have provided researchers with a statistical tool capable of examining correlation among a full set (triplicate) of replicates within a single test, rather than multiple pair-wise comparisons. Removing co-zeros did little to improve the sensitivity of this statistic. The other statistic from Kendall, Kendall’s *τ*, displayed similar performance to the other non-parametric statistic, Spearman’s *ρ*. This finding is contrary to other studies of Kendall’s *τ* conducted in the fields of signal processing and psychology [43, 44]. For analysis of genomic data, the Spearman’s *ρ* is standard in deeptools’ correlation functions [13]

Of the statistics examined here, the *R*^2^ coefficient and normalized mutual information score were the most sensitive to the change in shared peaks between replicates (when co-zeros were removed). Comparison of these two statistics revealed that normalized mutual information was the better-behaved statistic. This behavior manifested as smaller AUC within simulations, less variation in values across simulations, and smoother values between unrelated synthetic replicates. Similarly, the computational evidence provided by our random forest model suggests that normalized mutual information was better at estimating experimental relationships between true ATAC-seq replicates. Taken together, these results indicate that of the two metrics, normalized mutual information may be the stronger association metric for ATAC-seq data. Information theoretic approaches, such as normalized mutual information, have been utilized in several other biological fields, ranging from cancer genomics to fungal genetics [51, 52, 53, 54, 55, 56, 57]. Regarding ATAC-seq data, a handful of other studies have specifically used mutual information in data integration, analysis, and deep-learning of single-cell ATAC-seq profiles [58, 59]. For those investigator interested in using information theoretic approaches, several of these functions are made easily available within the python, scikit learn library [46].

Sparsity and zero mapped sequenced reads are not unique properties of ATAC-seq data. These extend to genomic, Hi-C, ChIP-seq, and RNA-seq data sets. Imputation along with modified zero-inflated models have been used with success for studying RNA sequencing data sets and detecting regions with differential expression [60]. Simulations and models of sampling zero-genomic count data have been developed to understand the effects of these values, particularly in the context of differential analysis [36]. Previous simulation studies of ATAC-seq have been focused on generating ATAC-seq data, for pipeline development, or single-cell ATAC-seq samples, to examined different approaches in their analysis [61, 62]. To our knowledge, this is the first example of using a simulation approach for studying reproducibility and association of ATAC-seq samples. Adapting strategies from these previous studies will help improve our simulation and expand it to other genomic and epigenomic sequencing data. The current results of our study strongly suggest that normalized mutual information is an appropriate metric for measuring reproducibility in chromatin accessibility assays.

## Conclusions

For this study, we produced eight ATAC-seq experiments using the A549 Cancer cell line. Across replicates, these ATAC-seq samples are well correlated and reproducible. For investigations of chromatin accessibility (particularly in the A549 cell line), these experiments are an additional resource for developing analysis pipelines, peak detection algorithms, and machine learning approaches.

Leveraging the A549 ATAC-seq experiments, we designed a computational simulation to generate simulated replicates. Specifically, synthetic replicates were coded that share a known, fixed portion of significantly enriched loci. Using these replicates, correlation metrics—the Pearson’s *R*, Spearman’s *ρ*, Top-Down, and Kendall’s *τ* —and association statistics (ranging from zero to one—the *R*^2^ coefficient, Kendall’s W, and normalized mutual information—were tested for accuracy. Overall, the reported value of these statistics was inflated and much larger than the fixed portion of shared, significant loci between replicates.

Removing specific outliers from ATAC-seq data, specifically the removal of co-zeros, improved estimates of correlation and association. We estimate that co-zero values, when comparing WFpkm counts between two real ATAC-seq experiments, occupy nearly five percent of a bi-variate distribution. While only a small portion of the total data, filtering these values from analysis greatly improves the measurements of most correlation and association statistics between samples, in simulation. Applied to real ATAC-seq data, removing co-zero values from comparison significantly reduced the reported correlation and association statistic, matching results from simulation.

One of the association statistics examined here is normalized mutual information, an information theoretic approach that is less well known across the (epi)genomics field. After removing co-zero values, normalized mutual information displayed the lowest inflation relative to the similarity between simulated replicates. The *R*^2^ coefficient also performed well in simulations (after removal of co-zeros), displaying good sensitivity to differences between simulated replicates. Of these two association metrics, a random forest model selected normalized mutual information as the stronger feature when estimating experimental relationships between real ATAC-seq experiments. From these results we conclude that normalized mutual information is a powerful, non-parametric approach for estimating association between ATAC-seq experiments.

## Methods

### Construction of A549 ATAC-seq libraries

ATAC-seq experimental libraries were generated using A549 human lung carcinoma epithelial cells (ATCC, VA, catalog #CCL-185) [63, 64, 65]. Three biological replicate libraries were prepared from freshly harvested cells using an ATAC-seq kit (Active Motif, 53150) following the manufacturer’s protocol. The remaining five libraries were prepared using cryopreserved cells following methods outlined in Milani *et al.* (2016) with modifications [18]. Briefly, A549 cells were cultured in T75 flasks and harvested by trypsinization using 0.25% (w/v) Trypsin-EDTA (0.5%) solution (Gibco, 15400054). Harvested cells were centrifuged and pellets resuspended in freezing media containing DMEM (Gibco, 11885-084), 10% FBS (Corning, 35-015-CV), and 10% DMSO (ATCC, 4-X). Pellets were frozen using an isopropyl alcohol chamber (Thermo Fisher Scientific, 5100-0001) at –80*^◦^*C. After 24 hours, frozen cells were transferred to liquid nitrogen for long term storage. To perform experiments, cryopreserved cells were transferred to –80*^◦^*C for several days, and the tube was immersed in 37°C water bath for approximately two minutes on the day libraries were prepared. Thawed cells were resuspended in 1X PBS with protease inhibitor cocktail (Thermo Fisher Scientific, 78430). Cell counts and viability were assessed and aliquots containing 80,000 cells per sample were processed into ATAC-seq libraries.

### Sequencing, alignment and filtering

ATAC-seq libraries were sequenced at the sequencing facility at Los Alamos National Laboratory on an Illumina NextSeq2000 sequencer in paired end mode (PE151) using P3 chemistry. With Fastp, raw reads were trimmed and filtered to remove Nextra adaptors and reads with repetitive sequences [66]. Additionally reads were also filtered to remove bases with low quality scores (q *<* 15). These processed reads were aligned to the new, telomere-to-telomere human reference genome, version 2 [67] via bwa [68]. After alignment, duplicate sequenced pairs were marked via samblaster and removed from analysis [69]. Read pairs mapping to the mitochondria were also removed (see Supplementary Table S1).

### Other data used

Raw ATAC-seq data, in the form of paired fastq.gz files, was downloaded from the ENCODE project for the A549, HepG2, RWPE2, GM12878, IMR-90, K562, and WTC11 cell lines [70, 11]. The ENCODE file experiment and replicate accession numbers are included in Table 1. For alignment, these data were passed through the same pipeline described above for ATAC-seq samples generated here, and aligned to the human, telomere-to-telomere, reference genome [67].

### Peak calling, peak filtering and reproducibility

After filtering, sample alignments were analyzed to identify loci displaying significant enrichment of paired-end reads. This peak calling was conducted using MACS2 [6, 71]. Specifically, after removing duplicates and mitochondrial mapped reads, samples were further filtered using samtools with the following flags: -F 4 -F 256 -F 512 -F 1024 -F 2048 -q 30 and then passed to MACS2 in BAMPE mode [72, 73].

Between true, biological replicates, reproducible peaks were identified via irreproducible discovery rate thresholding [74]. Using ChIP-R, replicate narrow peak files were filtered to retain only those peaks that were consistent across all replicates; in ChIP-R, where command line parameter, m = number of biological replicates [75]. In addition to this setting the ’-fragment’ option was also invoked. These sets of final peak counts were retained for further analysis.

### Genomic down-sampling and simulation design

For each of the eight ATAC-seq experiments of A549 cells generated in this study, synthetic replicates were generated by duplicating a given sample into two copies and then randomly, varying the total number of shared peaks between them. Specifically, for a given ATAC-seq experiment, a set portion of peaks was chosen at random, such that within one of the synthetic replicates, a given selected peak was depleted, randomly removing a portion of the alignments within the peak bounds (as defined by MACS2). These sets of peaks were randomly selected from the set of reproducible peaks for that sample and its associated biological replicates (see above). Three sets of simulations were conducted, removing 50, 85 and 95% of reads within selected peaks. This procedure results in two synthetic ATAC-seq replicates, generated from a single, true parent ATAC-seq data set. These synthetic ‘sister’ ATAC-seq data sets have identical genome-wide alignments except within a sub-set of loci that vary between them. From each true ATAC-seq data set, synthetic sister replicates were generated by varying the total percentage of shared peaks from 99 to 5%, with a delta of 5%. For each simulation, across the change in portion of shared peaks, a common random seed was used to preserve autocorrelation across this axis. This process was repeated fifteen times for each of the eight, A549 ATAC-seq samples, totaling a one hundred and twenty simulations.

### Genomic binning, fragment counts, and standardization

On both synthetic samples from simulation studies or replicates from (true) ATAC-seq experiments, a genomic binning approach was used to estimate correlation and association statistics between samples. For each chromosome, contiguous bins were established 5’–3’, every ten kilobases. Within each of these bins, the number of sequenced fragments is counted and standardized to fragments per kilobase per million. These counts were rounded up to their nearest whole integer generating standardized counts of whole fragments per kilobase per million (WFpkm).

### Calculating correlation and association metrics

In python scripts, using the scipy-stats module [76], the Pearson’s *R*, Spearman’s *ρ*, and Kendall’s *τ* were calculated on the WFpkm counts between pairs of ATAC-seq replicates. Functions for the Top-Down correlation metric [48] and Kendall’s W rank statistic [50, 41] were also developed using custom python code. The *R*^2^ coefficient was calculated using the square of the Pearson’s *R*. The normalized mutual information statistic from pythons sklearn module [46] was used in association studies. Between any pair of WFpkm counts, the bi-variate distribution was examined to identify instances were both profiles contained a value of zero WFpkm. For studies of the effects of co-zero inflation, these co-zero values were removed, and the correlation (or association) statistics recalculated on these filtered distributions.

For correlation analysis on ATAC-seq experiments conducted here using A549 cells, the Pearson’s *R* correlation statistic was calculated on WFpkm values between replicates with co-zeros removed. Similarly, co-zeros were removed prior to calculating correlation and association statistics between replicates of ATAC-seq data downloaded from the ENCODE project public repository.

### Statistical tests on area under the curve

Across simulations, values of correlation and associations statistics were calculated as a function of the designed portion of peaks between synthetic replicates. For each statistic tested, the 95% confidence interval of the average area under the curve was calculated via bootstrapping, with a thousand iterations. This was done for statistical profiles from simulations with and without co-zero values. For comparisons of the average area under the curve, either between statistics or within the same statistic after removing co-zeros, one thousand permutations were used to calculate the null distribution of the difference between the mean area under the curve [77]. The proportion of these differences greater than or equal to the true observed difference was used as the *p*-value. A significance level of 0.05 was used to reject the null hypothesis, H_0_: no difference in mean area under the curve, in favor of our alternative hypothesis, H_1_: difference of mean area under the curve.

### Design of random forest model

A random forest model was built in python using the scikit learn module [47, 46]. Association statistics from the ATAC-seq data generated in this study on A549 cells and additional ATAC-seq data downloaded from the ENCODE project was used as input (see Table 1). As features in this random forest, the *R*^2^ coefficient and normalized mutual information were calculated between every pair of ATAC-seq experiments using WFpkm counts, across ten kilobase pair, genomic bins and removing co-zero values. The comparison of each unique pair of experiments (totaling 276) were discretized as (1) between independent ATAC-seq experiments in different cell lines, (2) independent experiments using the same cell line, and (3) between true replicates. The total number of comparisons distributed among these three classes was 213, 45, and 18 (respectively). Given the over-representation of comparison between independent ATAC-seq experiments in different cell lines, 39 of the 213 comparisons were chosen randomly to represent the total, unique comparisons between experiments with unique cell lines. This down sampling resulted in 39, 45, and 18 comparisons between independent experiments in different cell lines, independent experiments using the same cell line, and true replicate experiments, respectively.

For the testing and training of the model, test and training sets of the classes defined above were selected using a stratified, 40:60 split of the data. Additionally, ten-fold, stratified cross validation was used to train and test the model [78]. A hundred estimators with the entropy selection criterion were used along with default settings in the python random forest classifier function within scikit learn [46].

## Declarations

### Ethics approval and consent to participate

Not applicable

### Consent for publication

Approved for public release; distribution is unlimited: LA-UR-23-24317

### Data availability

All data and code associated with this manuscript (if not publicly available) is available upon request. Raw sequence reads generated by this study on A549 samples are deposited and stored on NCBI’s Sequence Read Archive, with accession numbers SAMN35335737 – SAMN35335744. Scripts, code, and software used in the statistical analysis and visualization are stored on GitHub: https://github.com/cjr41/SLURPY/tree/ main/ATAC_SEQ_SIMULATION

### Authors’ contributions

The authors (with initials) CR, VV, VJ, NL, KYS, CRS, and SRS contributed to the overall experimental design of this manuscript. VV and CRS provided materials and wrote experimental methods. VV prepared experimental ATAC-seq data for cultured A549 cells. CR conducted analysis and produced visualizations. CR, CRS, and SRS wrote the paper. All authors edited and provided comments on the text of the manuscript.

## Acknowledgements

We would like to thank members of the Genomics and Bioanalytics Group at Los Alamos National Laboratory for their helpful comments on this manuscript. Special thanks to Drs. Taehyung Kwon and Colin Kruse for their specific and helpful suggestions on analyses. We would also like to acknowledge and thank the ENCODE Consortium and the members of the Michael Snyder lab of Stanford, for generating the ATAC-seq data sets used within this study.

## Supplementary Materials

### Additional File 1

- Title: Statistical profiles Simulation 0.50.png
- File Format: png
- Description: Correlation and association values (y-axis) as a function of percentage of shared peaks between synthetic replicates (x-axis). Red and grey curves depict the mean and 95% CI (respectively) values across simulations. A grey, dashed line marks a one-to-one relationship between the x- and y-axis. Left and right columns display change in values as a function of removing co-zeros. Results are from simulations with 50% paired reads within selected peaks removed.

### Additional File 2

- Title: Statistical profiles Simulation 0.95.png
- File Format: png
- Description: Correlation and association values (y-axis) as a function of percentage of shared peaks between synthetic replicates (x-axis). Red and grey curves depict the mean and 95% CI (respectively) values across simulations. A grey, dashed line marks a one-to-one relationship between the x- and y-axis. Left and right columns display change in values as a function of removing co-zeros. Results are from simulations with 95% paired reads within selected peaks removed.

---

**Figure S1:**
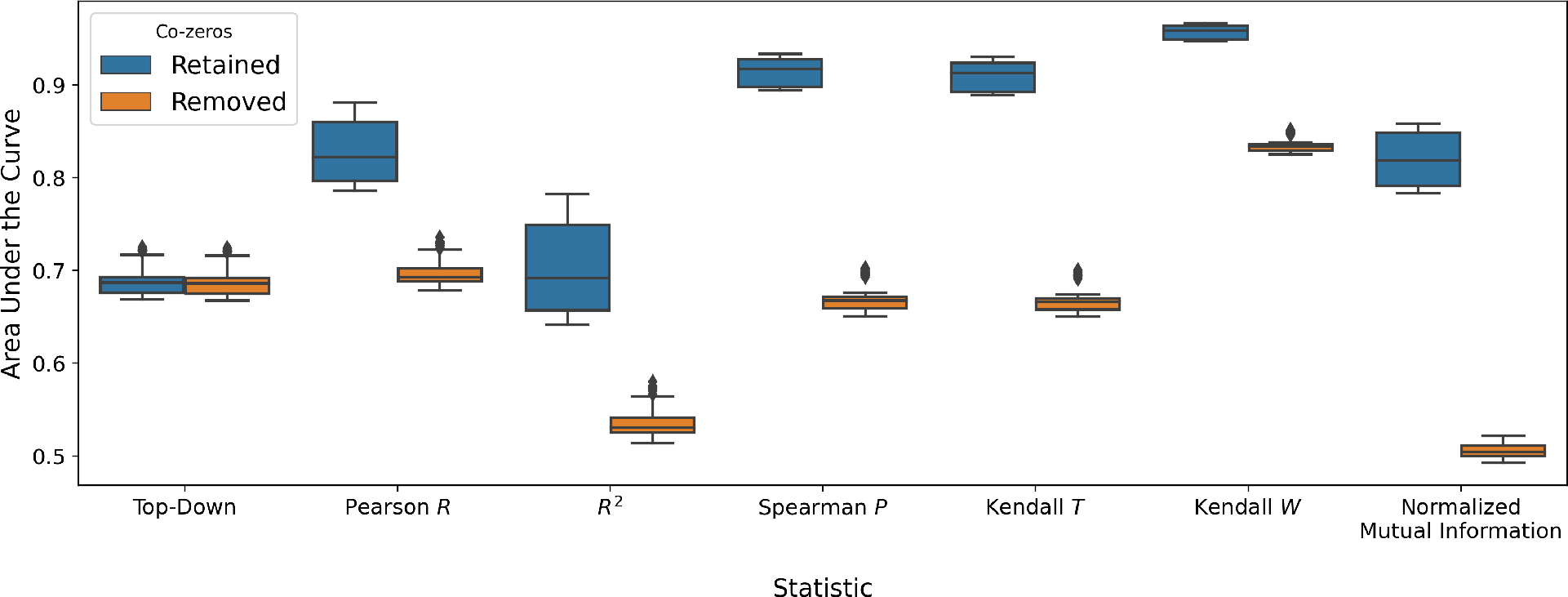
Boxplots displaying the area under the curve (y-axis) across statistics (x-axis) with co-zeros retained and removed from analysis (blue and orange boxes, respectively).

**Figure S2:**
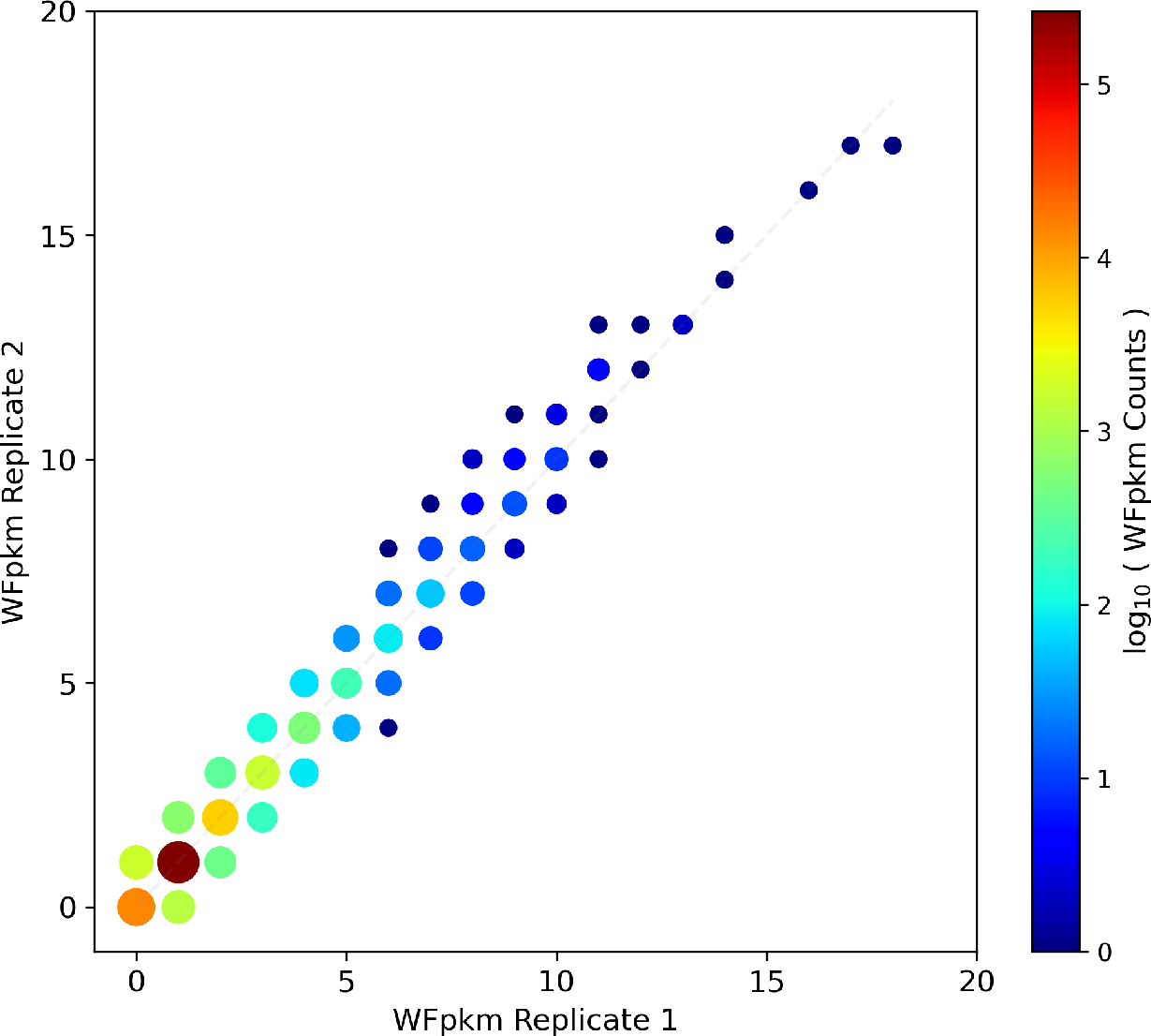
Bi-variate plot of WFpkm counts (across 10 kb genomic bins) between replicates of real, A549 ATAC-seq experiments. Dark red to blue colors and marker size designate the density (log10 (WFpkm counts)) of counts between replicates. Co-zero values appear as an orange dot in lower left corner. A dashed grey line represents a one-to-one relationship between the two replicates.

**Figure S3:**
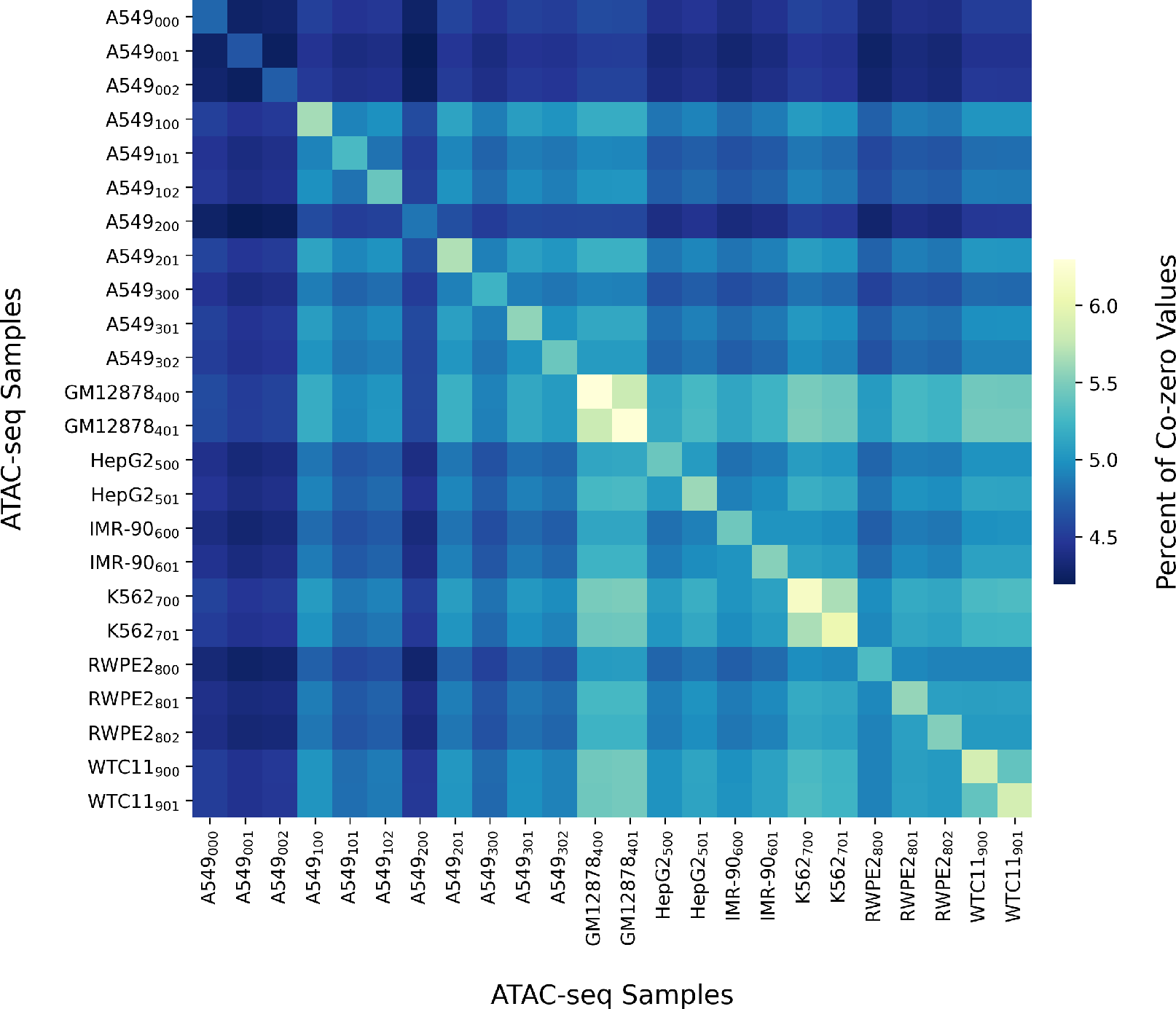
The percent of co-zero values in bi-variate WFpkm distributions between real ATAC-seq experiments. Sample names are annotated along the x- and y-axis.

**Figure S4:**
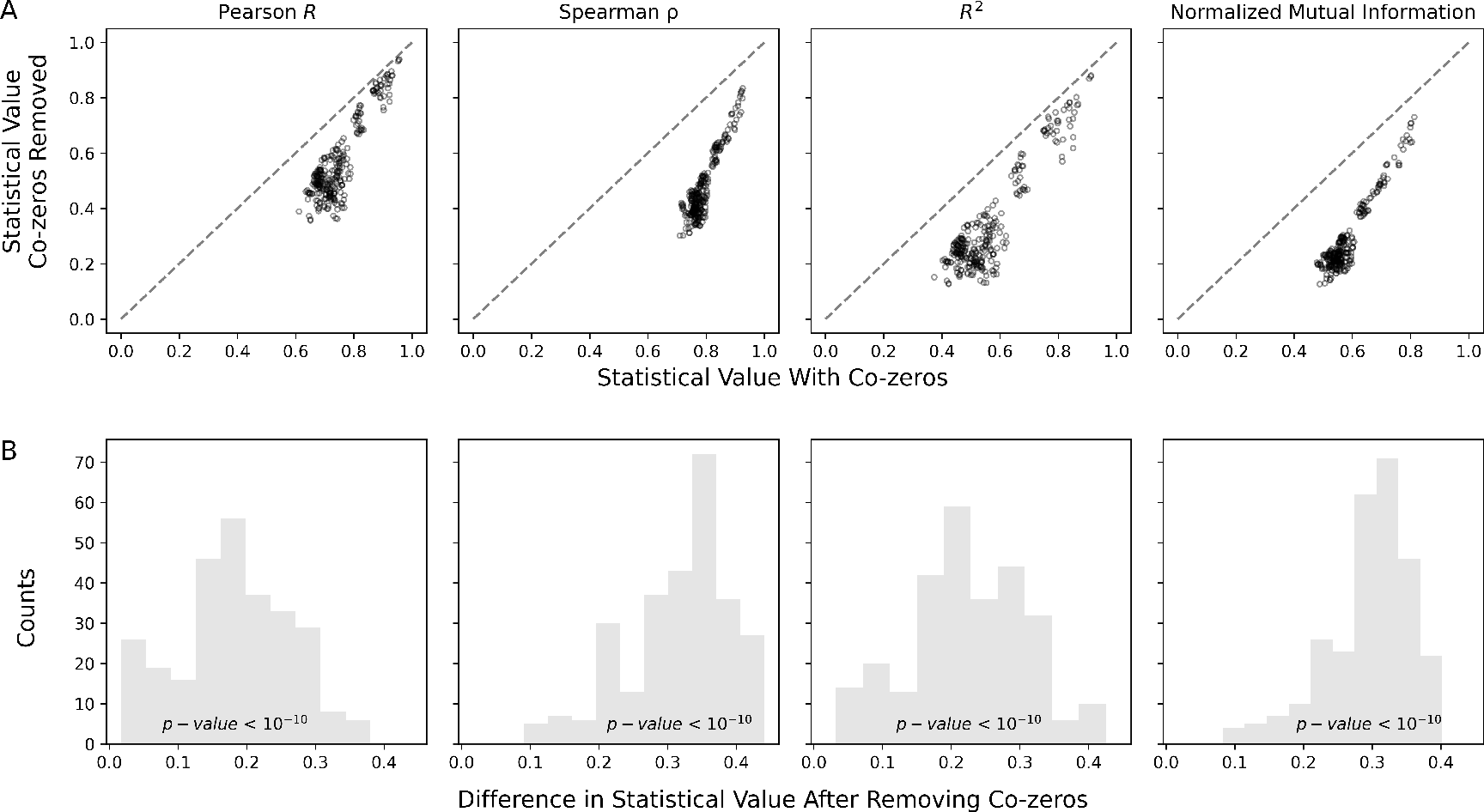
The effect of removing co-zeros from analysis on ATAC-seq experiments from the ENCODE project. **A** Shift in the estimates of correlation and association before (x-axis) and after (y-axis) removing co-zeros from analysis for the Pearson’s *R*, the Spearman’s *ρ*, *R*^2^ coefficient, and normalized mutual information, left to right respectively. A dashed line denotes a one-to-one relationship. **B** The pair-wise difference in correlation and association metrics from estimates before and after removing co-zeros.

**Figure S5:**
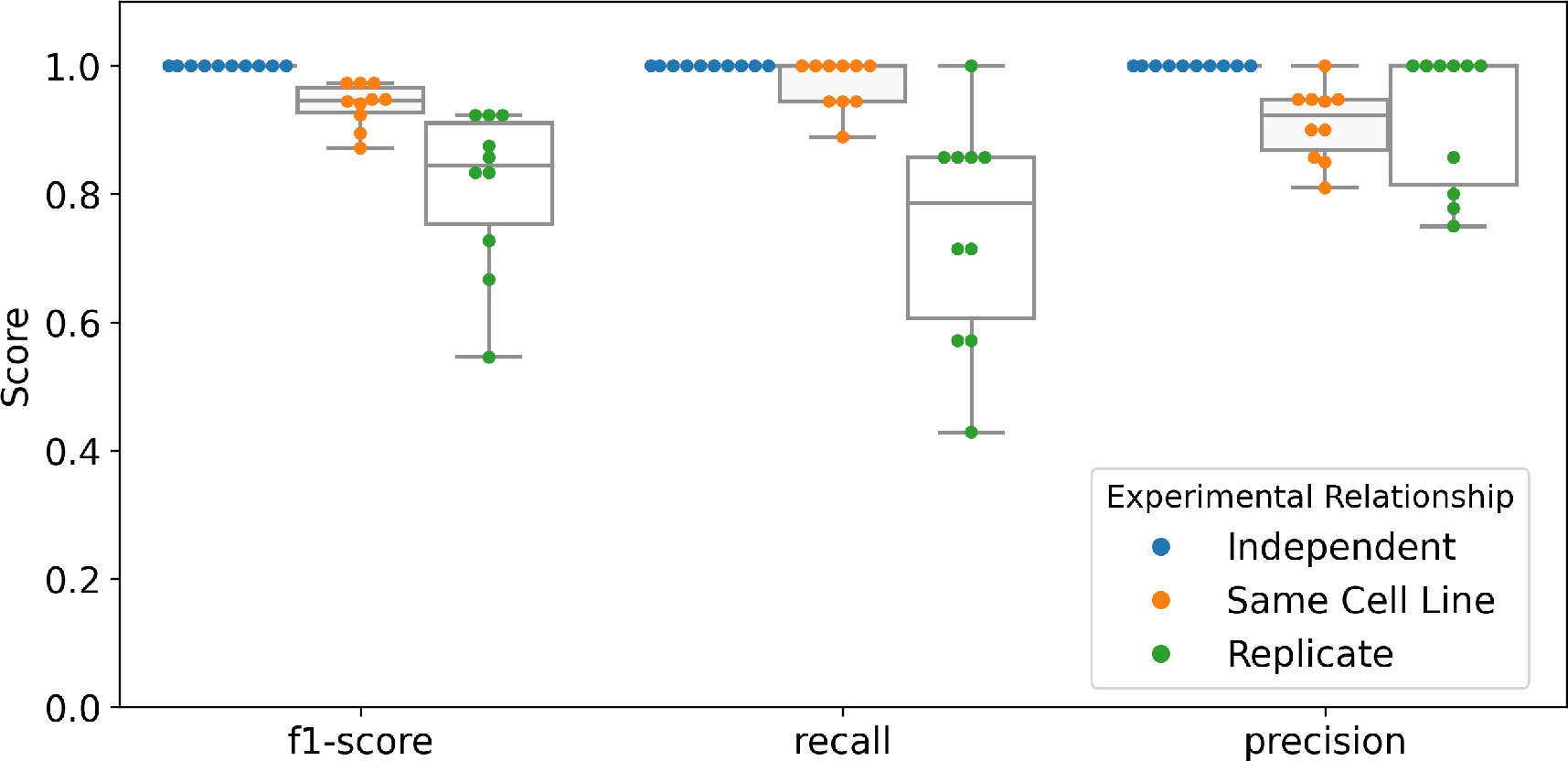
The f1-scores, recall, and precision of the random forest model with ten-fold, stratified cross validation. Blue, orange, and green colors denote experimental relationship class.

**Table S1:** Read Counts of ATAC-seq Experiments

| Sample Title | Cell Line | Total Reads | Mapped Reads | Not Used | mtDNA | Duplicates | Un-mapped | Low Quality Reads | Replicate Name | Source |
| --- | --- | --- | --- | --- | --- | --- | --- | --- | --- | --- |
| A549<sub>000</sub> | A549 | 341325836 | 259029456 | 21246814 | 12009324 | 46948944 | 408486 | 1682812 | ENCLB404SKN | ENCSR032RGS |
| A549<sub>001</sub> | A549 | 442074976 | 329679445 | 27536117 | 15475857 | 66338506 | 506856 | 2538195 | ENCLB605LCC | ENCSR032RGS |
| A549<sub>002</sub> | A549 | 277970512 | 211291691 | 18456829 | 11170486 | 35323112 | 343051 | 1385343 | ENCLB817BKI | ENCSR032RGS |
| A549<sub>100</sub> | A549 | 65405524 | 23987725 | 2973344 | 33653170 | 3093813 | 48906 | 1648566 | 2501_001 | This study |
| A549<sub>101</sub> | A549 | 84816540 | 22605005 | 2595465 | 55231224 | 2481489 | 32350 | 1871007 | 2501_002 | This study |
| A549<sub>102</sub> | A549 | 64084756 | 17618743 | 2122339 | 40809830 | 1979951 | 67472 | 1486421 | 2501_003 | This study |
| A549<sub>200</sub> | A549 | 133625408 | 35069198 | 4386111 | 86279921 | 4418826 | 31955 | 3439397 | 2501_007 | This study |
| A549<sub>201</sub> | A549 | 69273610 | 15377297 | 1834370 | 48556785 | 1780806 | 27699 | 1696653 | 2501_008 | This study |
| A549<sub>300</sub> | A549 | 86963986 | 42567716 | 4788405 | 31620108 | 5703777 | 121028 | 2162952 | 2501_018 | This study |
| A549<sub>301</sub> | A549 | 84297712 | 28744542 | 3582876 | 44737775 | 5400961 | 118620 | 1712938 | 2501_019 | This study |
| A549<sub>302</sub> | A549 | 97877188 | 35836016 | 4491769 | 49816243 | 5663997 | 42546 | 2026617 | 2501_020 | This study |
| GM12878<sub>400</sub> | GM12878 | 76479882 | 46889870 | 4260513 | 11245729 | 12635046 | 252057 | 1196667 | ENCLB584REF | ENCSR095QNB |
| GM12878<sub>401</sub> | GM12878 | 69456510 | 49588811 | 4319318 | 7334534 | 6740176 | 186878 | 1286793 | ENCLB907YRF | ENCSR095QNB |
| HepG2<sub>500</sub> | HepG2 | 76077306 | 48113686 | 6037783 | 8348893 | 11668633 | 235020 | 1673291 | ENCLB074EQT | ENCSR042AWH |
| HepG2<sub>501</sub> | HepG2 | 88838406 | 48246610 | 6580203 | 19207768 | 12021756 | 605060 | 2177009 | ENCLB324GIU | ENCSR042AWH |
| IMR-90<sub>600</sub> | IMR-90 | 84117916 | 47543633 | 11830808 | 8448694 | 9559287 | 5990188 | 745306 | ENCLB432QLN | ENCSR200OML |
| IMR-90<sub>601</sub> | IMR-90 | 95034796 | 61359070 | 6202820 | 14378540 | 10233756 | 1872742 | 987868 | ENCLB937FOM | ENCSR200OML |
| K562<sub>700</sub> | K562 | 78745422 | 48217636 | 6777147 | 10759718 | 10705486 | 91659 | 2193776 | ENCLB758GEG | ENCSR483RKN |
| K562<sub>701</sub> | K562 | 83982064 | 52270533 | 6752478 | 10447009 | 12175330 | 162811 | 2173903 | ENCLB918NXF | ENCSR483RKN |
| RWPE2<sub>800</sub> | RWPE2 | 67263926 | 55152003 | 6718663 | 753542 | 2685741 | 286519 | 1667458 | ENCLB293SLX | ENCSR080SNF |
| RWPE2<sub>801</sub> | RWPE2 | 53441754 | 43166947 | 5244472 | 1277887 | 2088775 | 323207 | 1340466 | ENCLB734LAL | ENCSR080SNF |
| RWPE2<sub>802</sub> | RWPE2 | 60212304 | 48162285 | 5946340 | 1888964 | 2288274 | 331284 | 1595157 | ENCLB984XHJ | ENCSR080SNF |
| WTC11<sub>900</sub> | WTC11 | 115952320 | 74558506 | 7218516 | 7595855 | 21538753 | 4422396 | 618294 | ENCLB621FEI | ENCSR541KFY |
| WTC11<sub>901</sub> | WTC11 | 127343084 | 79335328 | 7889715 | 9553155 | 24757738 | 5028262 | 778886 | ENCLB715JYV | ENCSR541KFY |

